## Supporting Information for "Repeat DNA-PAINT suppresses background and non-specific signals in optical nanoscopy"

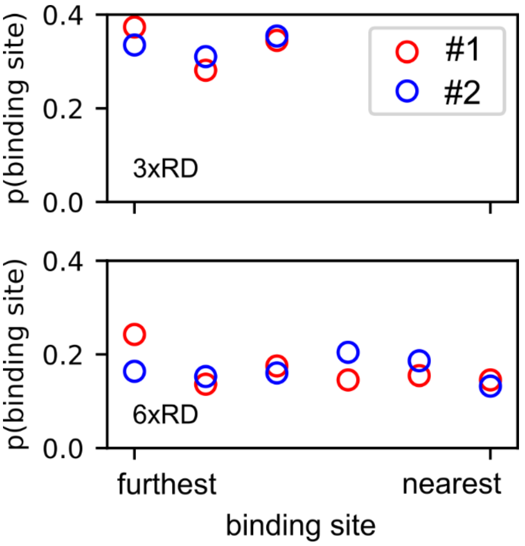


**Supplementary Figure 1.** Probability of the imager binding to each of the repeat binding domains in 3xRD (top) and 6xRD (bottom) docking motifs as determined from Forward-Flux Sampling simulations carried out with oxDNA. See Online Methods for details. The probabilities are approximately equal between domains. Red and blue points correspond to two independent computational runs.

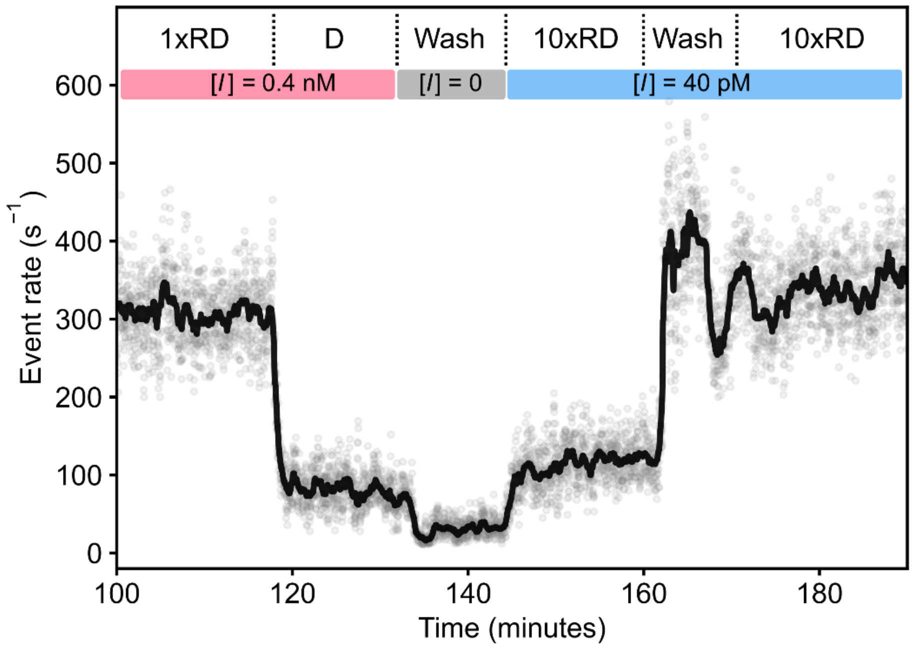


**Supplementary Figure 2.** A partial time trace of event-rates over time recorded in cardiac tissue where RyRs are labelled with the DNA constructs in **Fig. 1c**. Data are shown starting from 100 minutes after the beginning of the experiment, whilst imaging 1xRD at high imager concentration, [*I*] = 0.4 nM. When the Displacer strand (D) was added with the imager still present, event rates dropped to typical background levels produced by non-specific binding. The following washing step further reduced event rate by removing leftover imager, along with D-1xRD duplexes (**Fig. 1c**). The 10xRD motif was then added along with the imager at a [*I*] = 40 pM, and a modest increase in event rate was observed. At this stage, event rates do not reach the same level as in the first experimental phase, due to excess 10xRD motifs that sequestrate available P1 in solution. After allowing 15 minutes for the hybridization between 10xRD and anchors strands to complete, excess 10xRD was washed away while retaining [*I*] = 40 pM, to restore event rates similar to those recorded in the first experimental phase. Note that all events are included in the event rate calculation, including non-specific events. As we show in Fig. 3, main text, the contribution of non-specific events is greater when using 0.4 nM imager (1xRD), ~8%, vs <1% when using 40 pM and 10xRD. When this is taken into account, the data above shows that specific event rates are about the same when using 1xRD with 0.4 nM versus 10xRD with 40 pM, within the precision that arises from stochastic variability.

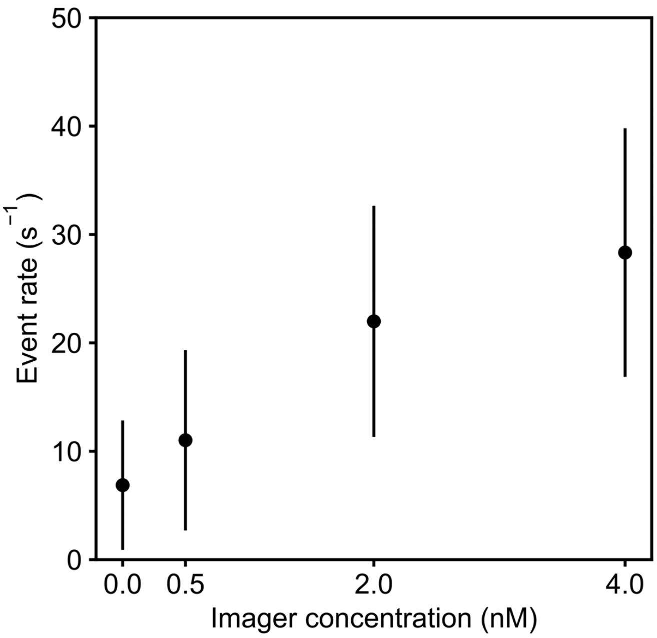


**Supplementary Figure 3.** Increase of non-specific event rate as a function of imager concentration in unlabeled cardiac tissue. The figure shows the dependency for the P5 imager labelled with ATTO 655 (**Supplementary Table 1**) whereas **Fig. 3a** shows the graph for imager P1. Qualitatively similar behavior is observed, indicating a near linear increase with imager concentration over a range of concentrations routinely used with DNA-PAINT. Error bars are calculated as mean ± SD from experimental repetitions.

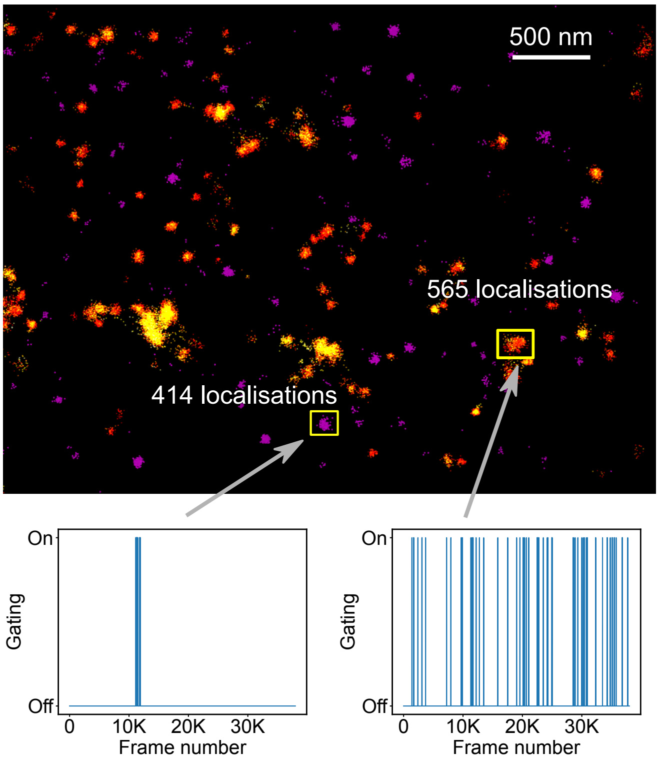


**Supplementary Fig. 4**. Non-specific P1 imager localizations (magenta, 5163 localizations) versus specific localizations (yellow-red, 33702 localizations) in a cardiac tissue sample in which RyRs are labelled. Boxes show a non-specific attachment (left) versus specific attachment (right) containing similar number of localizations. If the non-specific event occurred in a location close to areas of specific binding (see gating traces below) it would not be possible to detect and separate the non-specific events posthoc. This illustrates that non-specific imager binding at typical concentrations used in DNA-PAINT ([$I$]=0.4 nM) can significantly obscure and distort specific imager binding to docking strands with particular impact when targets are sparsely expressed. Note that we may not have detected all non-specific binding events in this image for just this reason, and thus underestimate the number of non-specific imager localizations. This limitation is greatly reduced with Repeat DNA-PAINT that reduces non-specific localizations ~10-fold while maintaining specific localizations at the same level.

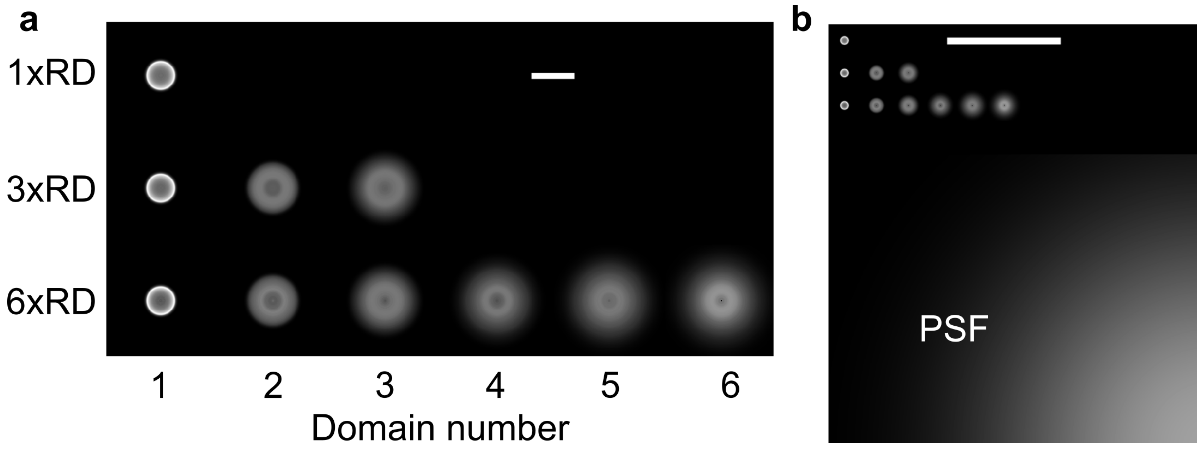


**Supplementary Figure 5. a**: 2D spatial projection of fluorophore location probability density for imagers hybridized to each of the binding domains of docking motifs 1xRD, 3xRD and 6xRD, as determined with coarse-grained molecular simulations (see Online Methods and **Supplementary Note 2**). These distributions are radially averaged around their center, corresponding to the docking motif anchoring point, to produce data in **Fig. 5a**. To aid the eye in comparing spot sizes, the brightness corresponding to a given probability density is not consistent between spots. Domains are increasingly numbered from the closest to the furthest from the anchoring point. Scale bar: 10 nm. **b:** Comparison between the distributions in panel **a** and the microscope PSF, modeled as an Airy disk with a full width half maximum (FWHM) of 250 nm. The Airy disk and fluorophore distributions are convolved and radially averaged to produce the curves in **Fig. 5b**. Scale bar: 100 nm.

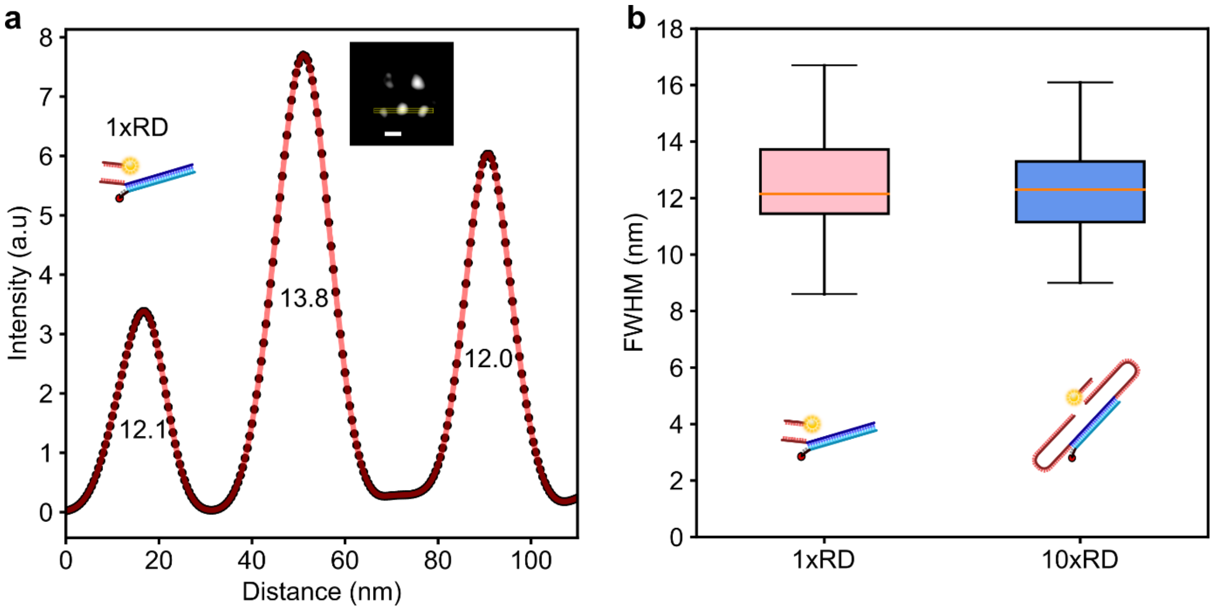


**Supplementary Figure 6. a:** Line intensity profile as recorded across the rendered image of an origami tile (see inset) with sites hosting 1xRD docking motifs. The width of the peaks is similar to that recorded with 10xRD motifs (**Fig. 5d**), demonstrating that Repeat DNA-PAINT preserves resolution. **b:** Distributions of the Full Width at Half Maximum (FWHM) of the peaks for both 1xRD (12.6 ± 2.1 nm) and 10x RD (12.3 ± 1.8 nm) motifs (mean ± SD) sampled over 30 individual sites across 10 tiles each, confirming that spatial resolution is unchanged between the two cases. Scale bar: 30 nm.

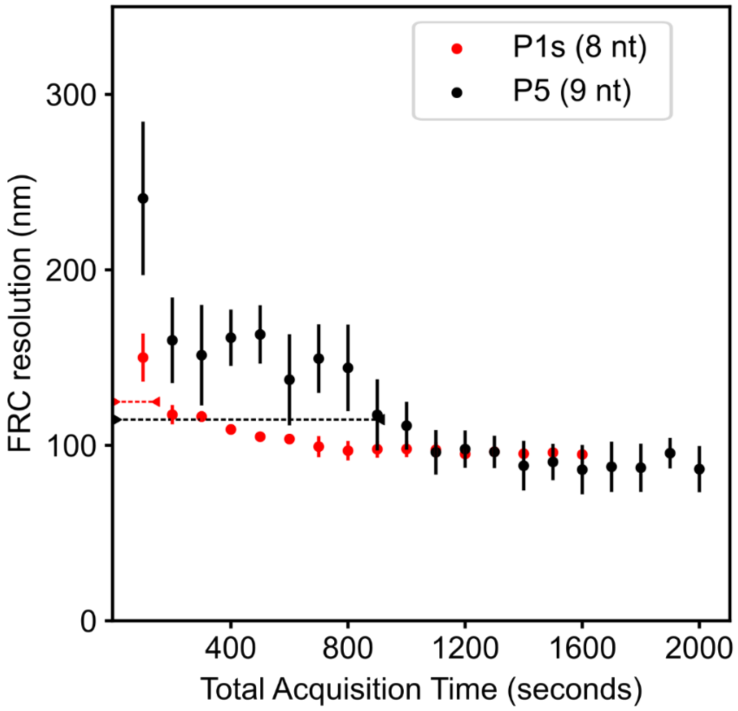


**Supplementary Figure 7.** Accelerated image acquisition with Repeat DNA-PAINT. With reference to the data presented in **Fig. 6b**, we compare the FRC image resolution over the course of two experiments, one conventional DNA-PAINT experiment employing 9 nt imager P5 (black) and one Repeat DNA-PAINT experiment where we use the shorter 8 nt imager P1s in combination with 10xRD docking motifs (red). Frame integration times were 100 ms for regular DNA-PAINT and 10 ms for Repeat DNA-PAINT, while the average duration of imager-docking binding events was ~40 ms for P5 and ~3 ms for P1s. In both experiments we used [*I*] ~0.3 nM. Measurement of FRC resolution versus total acquisition time indicates that a typical target resolution (~120 nm) was achieved about 6 times faster with Repeat DNA-PAINT. The fact that the imaging speed-up afforded by Repeat DNA-PAINT does not match the increase in frame-rate (10 fold) probably follows from the substantially lower number of photons recorded for P1s events compared to P5 events, leading to a lower localization precision for the Repeat DNA-PAINT run. This is however not an intrinsic limitation, as with a stronger laser source the photon yield of P1s could be increased to match those of P5, thus further accelerating Repeat DNA-PAINT imaging.

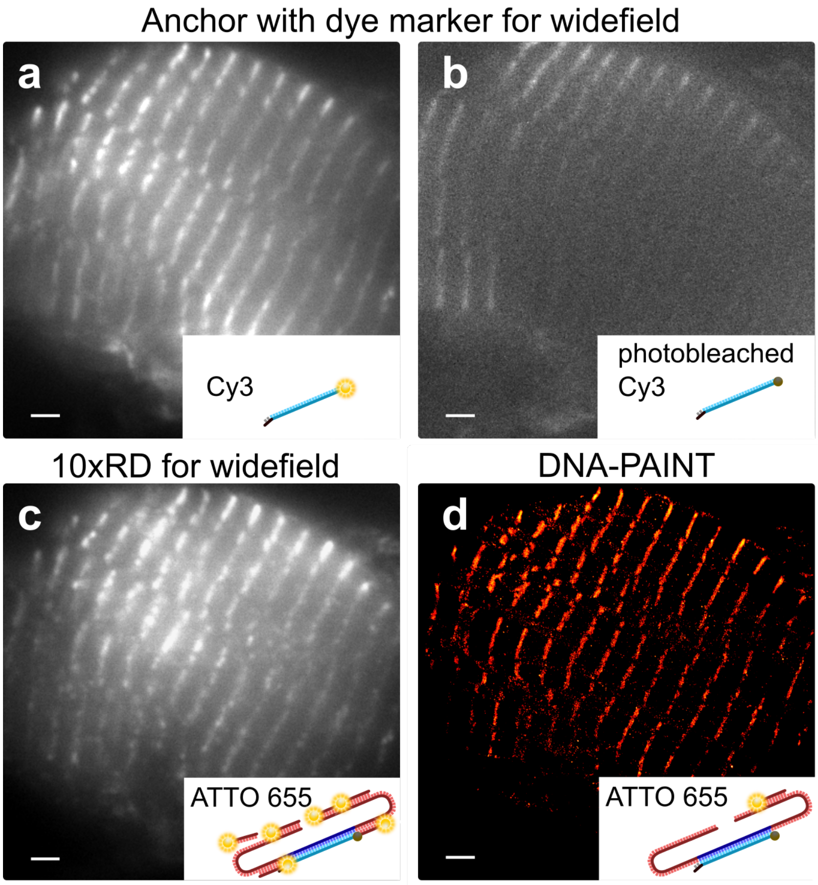


**Supplementary Figure 8.** Widefield functionality using repeat domains. **a:** Anchor strands with a 3' modified Cy3 fluorophore on tissue labelled via immunohistochemistry for alpha-actin were imaged. **b:** Not uncommon, after acquiring an image stack the Cy3 fluorophore shows signs of photobleaching. **c:** By functionalizing the anchor strand with 10x RD motifs an equivalent widefield image could be obtained using [$I$] = 1 nM of P1 ATTO 655 imager. **d:** Further reduction of imager concentration to 40 pM meant the tissue could then also be imaged as normal for super-resolution. Scale bar: 2 µm.

**Supplementary Table 1.** Sequences of the oligonucleotides used for DNA-PAINT measurements.

| Name | Sequence 5' → 3' |
|---|---|
| 1xRD (microsphere) | Biotin TT ATA CAT CTA |
| 3xRD (microsphere) | Biotin TT 3*{ATA CAT CTA} |
| 6xRD (microsphere) | Biotin TT 6*{ATA CAT CTA} |
| 1xRD (origami+tissue) | CTT CCT CAC AAT CAA AAT TTA CCT AAC ATA CAT CTA |
| 10xRD (origami+tissue) | 5*{ATA CAT CTA} CTT CCT CAC AAT CAA AAT TTA CCT AAC 5*{ATA CAT CTA} |
| Anchor (tissue) | Azide TTT TAG GTA AAT TTT GAT TGT GAG GAA G Cy5 |
| Anchor (origami) | TTT TAG GTA AAT TTT GAT TGT GAG GAA G |
| Displacer (D) | TAG ATG TAT GTT AGG TAA ATT TTG ATT GTG AGG |
| P1 (9bp) Imager | CTA GAT GTA T ATTO 655 |
| P5 (9bp) Imager | CTT TAC CTA A ATTO 655 |
| P1s (8bp) Imager | AGA TGT AT ATTO 655 |

**Supplementary Table 2.** Definitions of reaction coordinate used in the FFS simulations applied to calculate the relative binding rates of imagers to docking motifs (**Fig. 1b**). The minimum distance $d_{\mathrm{min}}$ and number of bonds $N_{\mathrm{bonds}}$ are both calculated over all pairs of nucleotides on the imager and docking strand (see Online Methods).

| $Q$ | Condition |
|---|---|
| -2 | $d_{\mathrm{min}} > 4$ |
| -1 | $3.5 < d_{\mathrm{min}} < 4$ |
| 0 | $1 < d_{\mathrm{min}} < 3.5$ |
| 1 | $N_{\mathrm{bonds}} = 0$ and $d_{\mathrm{min}} < 1$ |
| 2 | $1 < N_{\mathrm{bonds}} < 2$ |
| 3 | $2 < N_{\mathrm{bonds}} < 9$ |
| 4 | $N_{\mathrm{bonds}} = 9$ |

**Supplementary Table 3.** Number of simulated transitions sampled in FFS calculations to compute the initial flux $\Phi_{-2\to0}$ and the transition probabilities across the reaction-coordinate interfaces. For $\Phi_{-2\to0}$ the table shows the number of transitions sampled, while for the interface crossings we show the number of successful crossings and attempts (in brackets).

| | 1xRD Run 1 | 1xRD Run 2 | 3xRD Run 1 | 3xRD Run 2 | 6xRD Run 1 | 6xRD Run 2 |
|---|---|---|---|---|---|---|
| $\Phi_{-2\to0}$ | 15329 | 20068 | 16066 | 20083 | 20066 | 20075 |
| $\lambda_0^1$ | 100032 (710904) | 17086 (125539) | 71359 (461295) | 83376 (536040) | 100043 (603264) | 100034 (601579) |
| $\lambda_1^2$ | 20000 (1601182) | 20000 (1535431) | 20000 (1369312) | 20001 (1369327) | 20001 (1229732) | 20000 (1232036) |
| $\lambda_2^3$ | 20008 (120294) | 20015 (110998) | 20012 (105975) | 20009 (109635) | 20015 (99576) | 20087 (102229) |
| $\lambda_3^4$ | 20085 (109772) | 20083 (116510) | 20067 (90447) | 20080 (90953) | 20081 (84068) | 20087 (80362) |

**Supplementary Table 4.** Flux rates and success probabilities for each interface derived from the data in **Supplementary Table 3**. Since the order parameters for the three systems under study are very different, it may not be informative to compare success probabilities for a given interface between systems.

| | 1xRD Run 1 | 1xRD Run 2 | 3xRD Run 1 | 3xRD Run 2 | 6xRD Run 1 | 6xRD Run 2 |
|---|---|---|---|---|---|---|
| $\Phi_{-2\rightarrow 0}$ ($\times 10^6$ inverse timesteps) | 0.719 | 0.754 | 1.09 | 1.10 | 1.54 | 1.57 |
| $\lambda_0^1$ | 0.141 | 0.136 | 0.155 | 0.155 | 0.166 | 0.166 |
| $\lambda_1^2$ | 0.0125 | 0.0130 | 0.0146 | 0.0146 | 0.0163 | 0.0162 |
| $\lambda_2^3$ | 0.166 | 0.180 | 0.189 | 0.182 | 0.201 | 0.196 |
| $\lambda_3^4$ | 0.183 | 0.172 | 0.222 | 0.221 | 0.238 | 0.250 |
| Net flux ($\times 10^{11}$ inverse timesteps) | 0.385 | 0.413 | 1.03 | 1.00 | 1.99 | 2.07 |

**Supplementary Table 5.** DNA sequences of staples for DNA origami synthesis – staples with 5' biotin end modifications.

| **SEQUENCE** (5' → 3') |
|---|
| Biotin - ATTAAGTTTACCGAGCTCGAATTCGGGAAACCTGTCGTGC |
| Biotin - ATAAGGGAACCGGATATTCATTACGTCAGGACGTTGGGAA |
| Biotin - GCGATCGGCAATTCCACACAACAGGTGCCTAATGAGTG |
| Biotin - TTGTGTCGTGACGAGAAACACCAAATTTCAACTTTAAT |
| Biotin - ATTCATTTTTGTTTGGATTATACTAAGAAACCACCAGAAG |
| Biotin - CACCCTCAGAAACCATCGATAGCATTGAGCCATTTGGGAA |
| Biotin - AACAATAACGTAAAACAGAAATAAAAATCCTTTGCCCGAA |
| Biotin - AGCCACCACTGTAGCGCGTTTTCAAGGGAGGGAAGGTAAA |

**Supplementary Table 6**. DNA staple sequences for the origami designs used for quantifying event rates (**Fig. 1d**) and assessing imaging resolution (**Fig. 5c**). Anchor overhangs are emboldened.

| **SEQUENCE** (5' → 3') |
|---|
| TTTTCACTCAAAGGGCGAAAAACCATCACC |
| GTCGACTTCGGCCAACGCGCGGGGTTTTTC |
| TGCATCTTTCCCAGTCACGACGGCCTGCAG |
| TAATCAGCGGATTGACCGTAATCGTAACCG |
| AACGCAAAATCGATGAACGGTACCGGTTGA |
| AACAGTTTTGTACCAAAAACATTTTATTTC |
| TTTACCCCAACATGTTTTAAATTTCCATAT |
| TTTAGGACAAATGCTTTAAACAATCAGGTC |
| CATCAAGTAAAACGAACTAACGAGTTGAGA |
| AATACGTTTGAAAGAGGACAGACTGACCTT |
| AGGCTCCAGAGGCTTTGAGGACACGGGTAA |
| AGAAAGGAACAACTAAAGGAATTCAAAAAAA |
| CAAATCAAGTTTTTTGGGGTCGAAACGTGGA **TT AGG TAA ATT TTG ATT GTG AGG AAG** |
| CTCCAACGCAGTGAGACGGGCAACCAGCTGCA |
| TTAATGAACTAGAGGATCCCCGGGGGGTAACG |
| CCAGGGTTGCCAGTTTGAGGGGACCCGTGGGA |
| ACAAACGGAAAAGCCCCAAAAACACTGGAGCA |
| AACAAGAGGGATAAAAATTTTTAGCATAAAGC |
| TAAATCGGGATTCCCAATTCTGCGATATAATG |
| CTGTAGCTTGACTATTATAGTCAGTTCATTGA |
| ATCCCCCTATACCACATTCAACTAGAAAAATC |
| TACGTTAAAGTAATCTTGACAAGAACCGAACT **TT AGG TAA ATT TTG ATT GTG AGG AAG** |
| GACCAACTAATGCCACTACGAAGGGGGTAGCA **TT AGG TAA ATT TTG ATT GTG AGG AAG** |
| ACGGCTACAAAAGGAGCCTTTAATGTGAGAAT |
| AGCTGATTGCCCTTCAGAGTCCACTATTAAAGGGTGCCGT |
| GTATAAGCCAACCCGTCGGATTCTGACGACAGTATCGGCCGCAAGGCG |
| TATATTTTGTCATTGCCTGAGAGTGGAAGATT |
| GATTTAGTCAATAAAGCCTCAGAGAACCCTCA |
| CGGATTGCAGAGCTTAATTGCTGAAACGAGTA |
| ATGCAGATACATAACGGGAATCGTCATAAATAAAGCAAAG |
| TTTATCAGGACAGCATCGGAACGACACCAACCTAAAACGAGGTCAATC **TT AGG TAA ATT TTG ATT GTG AGG AAG** |
| ACAACTTTCAACAGTTTCAGCGGATGTATCGG **TT AGG TAA ATT TTG ATT GTG AGG AAG** |
| AAAGCACTAAATCGGAACCCTAATCCAGTT |
| TGGAACAACCGCCTGGCCCTGAGGCCCGCT |
| TTCCAGTCGTAATCATGGTCATAAAAGGGG |
| GATGTGCTTCAGGAAGATCGCACAATGTGA |
| GCGAGTAAAAATATTTAAATTGTTACAAAG |
| GCTATCAGAAATGCAATGCCTGAATTAGCA |
| AAATTAAGTTGACCATTAGATACTTTTGCG |
| GATGGCTTATCAAAAAGATTAAGAGCGTCC |
| AATACTGCCCAAAAGGAATTACGTGGCTCA |
| TTATACCACCAAATCAACGTAACGAACGAG |
| GCGCAGACAAGAGGCAAAAGAATCCCTCAG |
| CAGCGAAACTTGCTTTCGAGGTGTTGCTAA |
| AGCAAGCGTAGGGTTGAGTGTTGTAGGGAGCC |
| CTGTGTGATTGCGTTGCGCTCACTAGAGTTGC |
| GCTTTCCGATTACGCCAGCTGGCGGCTGTTTC |
| ATATTTTGGCTTTCATCAACATTATCCAGCCA |
| TAGGTAAACTATTTTTGAGAGATCAAACGTTA |

| |
|---|
| AATGGTCAACAGGCAAGGCAAAGAGTAATGTG |
| CGAAAGACTTTGATAAGAGGTCATATTTCGCA |
| TAAGAGCAAATGTTTAGACTGGATAGGAAGCC |
| TCATTCAGATGCGATTTTAAGAACAGGCATAG |
| ACACTCATCCATGTTACTTAGCCGAAAGCTGC |
| AAACAGCTTTTTGCGGGATCGTCAACACTAAA |
| TAAATGAATTTTCTGTATGGGATTAATTTCTT |
| CCCGATTTAGAGCTTGACGGGGAAAAAGAATA |
| GCCCGAGAGTCCACGCTGGTTTGCAGCTAACT |
| CACATTAAAATTGTTATCCGCTCATGCGGGCC |
| TCTTCGCTGCACCGCTTCTGGTGCGGCCTTCC |
| TGTAGCCATTAAAATTCGCATTAAATGCCGGA |
| GAGGGTAGGATTCAAAAGGGTGAGACATCCAA |
| TAAATCATATAACCTGTTTAGCTAACCTTTAA |
| TTGCTCCTTTCAAATATCGCGTTTGAGGGGGT |
| AATAGTAAACACTATCATAACCCTCATTGTGA |
| ATTACCTTTGAATAAGGCTTGCCCAAATCCGC |
| GACCTGCTCTTTGACCCCCAGCGAGGGAGTTA |
| AAGGCCGCTGATACCGATAGTTGCGACGTTAG |
| CCCAGCAGGCGAAAAATCCCTTATAAATCAAGCCGGCG |
| TAAATCAAAATAATTCGCGTCTCGGAAACCAGGCAAAGGGAAGG |
| GAGACAGCTAGCTGATAAATTAATTTTTGT |
| TTTGGGGATAGTAGTAGCATTAAAAGGCCG |
| GCTTCAATCAGGATTAGAGAGTTATTTTCA |
| CGTTTACCAGACGACAAAGAAGTTTTGCCATAATTCGA |
| TGACAACTCGCTGAGGCTTGCATTATACCAAGCGCGATGATAAA |
| TCTAAAGTTTTGTCGTCTTTCCAGCCGACAA |
| TCAATATCGAACCTCAAATATCAATTCCGAAA |
| GCAATTCACATATTCCTGATTATCAAAGTGTA |
| AGAAAACAAAGAAGATGATGAAACAGGCTGCG |
| ATCGCAAGTATGTAAATGCTGATGATAGGAAC |
| GTAATAAGTTAGGCAGAGGCATTTATGATATT |
| CCAATAGCTCATCGTAGGAATCATGGCATCAA |
| AGAGAGAAAAAAATGAAAATAGCAAGCAAACT |
| TTATTACGAAGAACTGGCATGATTGCGAGAGG |
| GCAAGGCCTCACCAGTAGCACCATGGGCTTGA |
| TTGACAGGCCACCACCAGAGCCGCGATTTGTA |
| TTAGGATTGGCTGAGACTCCTCAATAACCGAT |
| TCCACAGACAGCCCTCATAGTTAGCGTAACGA |
| AACGTGGCGAGAAAGGAAGGGAAACCAGTAA |
| TCGGCAAATCCTGTTTGATGGTGGACCCTCAA **TT AGG TAA ATT TTG ATT GTG AGG AAG** |
| AAGCCTGGTACGAGCCGGAAGCATAGATGATG |
| CAACTGTTGCGCCATTCGCCATTCAAACATCA |
| GCCATCAAGCTCATTTTTTAACCACAAATCCA |
| CAACCGTTTCAAATCACCATCAATTCGAGCCA |
| TTCTACTACGCGAGCTGAAAAGGTTACCGCGC |
| CCAACAGGAGCGAACCAGACCGGAGCCTTTAC |
| CTTTTGCAGATAAAAACCAAAATAAAGACTCC |
| GATGGTTTGAACGAGTAGTAAATTTACCATTA |
| TCATCGCCAACAAAGTACAACGGACGCCAGCA |
| ATATTCGGAACCATCGCCCACGCAGAGAAGGA |
| TAAAAGGGACATTCTGGCCAACAAAGCATC |
| ACCTTGCTTGGTCAGTTGGCAAAGAGCGGA |
| ATTATCATTCAATATAATCCTGACAATTAC |
| CTGAGCAAAAATTAATTACATTTTGGGTTA |
| TATAACTAACAAAGAACGCGAGAACGCCAA |
| CATGTAATAGAATATAAAGTACCAAGCCGT |

| |
|---|
| TTTTATTTAAGCAAATCAGATATTTTTTGT |
| TTAACGTCTAACATAAAAACAGGTAACGGA |
| ATACCCAACAGTATGTTAGCAAATTAGAGC |
| CAGCAAAAGGAAACGTCACCAATGAGCCGC |
| CACCAGAAAGGTTGAGGCAGGTCATGAAAG |
| TATTAAGAAGCGGGGTTTTGCTCGTAGCAT |
| TCAACAGTTGAAAGGAGCAAATGAAAAATCTAGAGATAGA |
| TCAAATATAACCTCCGGCTTAGGTAACAATTTCATTTGAAGGCGAATT |
| GTAAAGTAATCGCCATATTTAACAAAACTTTT |
| TATCCGGTCTCATCGAGAACAAGCGACAAAAG |
| TTAGACGGCCAAATAAGAAACGATAGAAGGCT |
| CGTAGAAAATACATACCGAGGAAACGCAATAAGAAGCGCA |
| GCGGATAACCTATTATTCTGAAACAGACGATTGGCCTTGAAGAGCCAC |
| TCACCAGTACAAACTACAACGCCTAGTACCAG |
| ACCCTTCTGACCTGAAAGCGTAAGACGCTGAG |
| AGCCAGCAATTGAGGAAGGTTATCATCATTTT |
| GCGGAACATCTGAATAATGGAAGGTACAAAAT |
| CGCGCAGATTACCTTTTTTAATGGGAGAGACT |
| ACCTTTTTATTTTAGTTAATTTCATAGGGCTT |
| AATTGAGAATTCTGTCCAGACGACTAAACCAA |
| GTACCGCAATTCTAAGAACGCGAGTATTATTT |
| ATCCCAATGAGAATTAACTGAACAGTTACCAG |
| AAGGAAACATAAAGGTGGCAACATTATCACCG |
| TCACCGACGCACCGTAATCAGTAGCAGAACCG |
| CCACCCTCTATTCACAAACAAATACCTGCCTA |
| TTTCGGAAGTGCCGTCGAGAGGGTGAGTTTCG |
| CTTTAGGGCCTGCAACAGTGCCAATACGTG |
| CTACCATAGTTTGAGTAACATTTAAAATAT |
| CATAAATCTTTGAATACCAAGTGTTAGAAC |
| CCTAAATCAAAATCATAGGTCTAAACAGTA |
| ACAACATGCCAACGCTCAACAGTCTTCTGA |
| GCGAACCTCCAAGAACGGGTATGACAATAA |
| AAAGTCACAAAATAAACAGCCAGCGTTTTA |
| AACGCAAAGATAGCCGAACAAACCCTGAAC |
| TCAAGTTTCATTAAAGGTGAATATAAAAGA |
| TTAAAGCCAGAGCCGCCACCCTCGACAGAA |
| GTATAGCAAACAGTTAATGCCCAATCCTCA |
| AGGAACCCATGTACCGTAACACTTGATATAA |
| GCACAGACAATATTTTTGAATGGGGTCAGTA |
| TTAACACCAGCACTAACAACTAATCGTTATTA |
| ATTTTAAAATCAAAATTATTTGCACGGATTCG |
| CCTGATTGCAATATATGTGAGTGATCAATAGT |
| GAATTTATTTAATGGTTTGAAATATTCTTACC |
| AGTATAAAGTTCAGCTAATGCAGATGTCTTTC |
| CTTATCATTCCCGACTTGCGGGAGCCTAATTT |
| GCCAGTTAGAGGGTAATTGAGCGCTTTAAGAA |
| AAGTAAGCAGACACCACGGAATAATATTGACG |
| GAAATTATTGCCTTTAGCGTCAGACCGGAACC **TT AGG TAA ATT TTG ATT GTG AGG AAG** |
| GCCTCCCTCAGAATGGAAAGCGCAGTAACAGT **TT AGG TAA ATT TTG ATT GTG AGG AAG** |
| GCCCGTATCCGGAATAGGTGTATCAGCCCAAT |
| AGATTAGAGCCGTCAAAAAACAGAGGTGAGGCCTATTAGT |
| GTGATAAAAAGACGCTGAGAAGAGATAACCTTGCTTCTGTTCGGGAGA |
| GTTTATCAATATGCGTTATACAAACCGACCGT |
| GCCTTAAACCAATCAATAATCGGCACGCGCCT |
| GAGAGATAGAGCGTCTTTCCAGAGGTTTTGAA |

| |
|---|
| GTTTATTTTGTCACAATCTTACCGAAGCCCTTTAATATCA |
| CAGGAGGTGGGGTCAGTGCCTTGAGTCTCTGAATTTACCGGGAACCAG **TT AGG TAA ATT TTG ATT GTG AGG AAG** |
| CCACCCTCATTTTCAGGGATAGCAACCGTACT **TT AGG TAA ATT TTG ATT GTG AGG AAG** |
| CTTTAATGCGCGAACTGATAGCCCCACCAG **TT AGG TAA ATT TTG ATT GTG AGG AAG** |
| CAGAAGATTAGATAATACATTTGTCGACAA |
| CTCGTATTAGAAATTGCGTAGATACAGTAC |
| CTTTTACAAAATCGTCGCTATTAGCGATAG |
| CTTAGATTTAAGGCGTTAAATAAAGCCTGT |
| TTAGTATCACAATAGATAAGTCCACGAGCA |
| TGTAGAAATCAAGATTAGTTGCTCTTACCA |
| ACGCTAACACCCACAAGAATTGAAAATAGC |
| AATAGCTATCAATAGAAAATTCAACATTCA |
| ACCGATTGTCGGCATTTTCGGTCATAATCA |
| AAATCACCTTCCAGTAAGCGTCAGTAATAA |
| GTTTTAACTTAGTACCGCCACCCAGAGCCA |

**Supplementary Table 7**. DNA staple sequences for the origami designs used for site loss (**Fig. 4**) and qPAINT measurements (**Fig. 6a**) (5' → 3'). Anchor overhangs targeted in experiment are (emboldened and colored red). Note that a second set of overhangs were present on this design (emboldened and colored blue) and could be directly probed by P1.

| **SEQUENCE** (5' → 3') |
|---|
| TTTTCACTCAAAGGGCGAAAAACCATCACC **TTA TAC ATC TAT TTC TTC ATT ATT CAC TTA CTA** |
| GTCGACTTCGGCCAACGCGCGGGGTTTTTC |
| TGCATCTTTCCCAGTCACGACGGCCTGCAG |
| TAATCAGCGGATTGACCGTAATCGTAACCG **TTT TAG GTA AAT T TTG ATT GTG AGG AAG** |
| AACGCAAAATCGATGAACGGTACCGGTTGA |
| AACAGTTTTGTACCAAAAACATTTTATTTC |
| TTTACCCCAACATGTTTTAAATTTCCATAT |
| TTTAGGACAAATGCTTTAAACAATCAGGTC |
| CATCAAGTAAAACGAACTAACGAGTTGAGA **TTT TAG GTA AAT T TTG ATT GTG AGG AAG** |
| AATACGTTTGAAAGAGGACAGACTGACCTT |
| AGGCTCCAGAGGCTTTGAGGACACGGGTAA |
| AGAAAGGAACAACTAAAGGAATTCAAAAAAA **TTA TAC ATC TAT TTC TTC ATT ATT CAC TTA CTA** |
| CAAATCAAGTTTTTTGGGGTCGAAACGTGGA |
| CTCCAACGCAGTGAGACGGGCAACCAGCTGCA |
| TTAATGAACTAGAGGATCCCCGGGGGGTAACG |
| CCAGGGTTGCCAGTTTGAGGGGACCCGTGGGA |
| ACAAACGGAAAAGCCCCAAAAACACTGGAGCA |
| AACAAGAGGGATAAAAATTTTTAGCATAAAGC |
| TAAATCGGGATTCCCAATTCTGCGATATAATG |
| CTGTAGCTTGACTATTATAGTCAGTTCATTGA |
| ATCCCCCTATACCACATTCAACTAGAAAAATC |
| TACGTTAAAGTAATCTTGACAAGAACCGAACT |
| GACCAACTAATGCCACTACGAAGGGGGTAGCA |
| ACGGCTACAAAAGGAGCCTTTAATGTGAGAAT |
| AGCTGATTGCCCTTCAGAGTCCACTATTAAAGGGTGCCGT |
| GTATAAGCCAACCCGTCGGATTCTGACGACAGTATCGGCCGCAAGGCG |
| TATATTTTGTCATTGCCTGAGAGTGGAAGATT |
| GATTTAGTCAATAAAGCCTCAGAGAACCCTCA |
| CGGATTGCAGAGCTTAATTGCTGAAACGAGTA |
| ATGCAGATACATAACGGGAATCGTCATAAATAAAGCAAAG |
| TTTATCAGGACAGCATCGGAACGACACCAACCTAAAACGAGGTCAATC |
| ACAACTTTCAACAGTTTCAGCGGATGTATCGG |
| AAAGCACTAAATCGGAACCCTAATCCAGTT |
| TGGAACAACCGCCTGGCCCTGAGGCCCGCT |
| TTCCAGTCGTAATCATGGTCATAAAAGGGG |
| GATGTGCTTCAGGAAGATCGCACAATGTGA |
| GCGAGTAAAAATATTTAAATTGTTACAAAG |
| GCTATCAGAAATGCAATGCCTGAATTAGCA |

| |
|---|
| AAATTAAGTTGACCATTAGATACTTTTGCG |
| GATGGCTTATCAAAAAGATTAAGAGCGTCC |
| AATACTGCCCAAAAGGAATTACGTGGCTCA |
| TTATACCACCAAATCAACGTAACGAACGAG |
| GCGCAGACAAGAGGCAAAAGAATCCCTCAG |
| CAGCGAAACTTGCTTTCGAGGTGTTGCTAA |
| AGCAAGCGTAGGGTTGAGTGTTGTAGGGAGCC |
| CTGTGTGATTGCGTTGCGCTCACTAGAGTTGC |
| GCTTTCCGATTACGCCAGCTGGCGGCTGTTTC |
| ATATTTTGGCTTTCATCAACATTATCCAGCCA |
| TAGGTAAACTATTTTTGAGAGATCAAACGTTA |
| AATGGTCAACAGGCAAGGCAAAGAGTAATGTG |
| CGAAAGACTTTGATAAGAGGTCATATTTCGCA |
| TAAGAGCAAATGTTTAGACTGGATAGGAAGCC |
| TCATTCAGATGCGATTTTAAGAACAGGCATAG |
| ACACTCATCCATGTTACTTAGCCGAAAGCTGC |
| AAACAGCTTTTTGCGGGATCGTCAACACTAAA |
| TAAATGAATTTTCTGTATGGGATTAATTTCTT |
| CCCGATTTAGAGCTTGACGGGGAAAAAGAATA |
| GCCCGAGAGTCCACGCTGGTTTGCAGCTAACT |
| CACATTAAAATTGTTATCCGCTCATGCGGGCC |
| TCTTCGCTGCACCGCTTCTGGTGCGGCCTTCC |
| TGTAGCCATTAAAATTCGCATTAAATGCCGGA |
| GAGGGTAGGATTCAAAAGGGTGAGACATCCAA |
| TAAATCATATAACCTGTTTAGCTAACCTTTAA |
| TTGCTCCTTTCAAATATCGCGTTTGAGGGGGT |
| AATAGTAAACACTATCATAACCCTCATTGTGA |
| ATTACCTTTGAATAAGGCTTGCCCAAATCCGC |
| GACCTGCTCTTTGACCCCCAGCGAGGGAGTTA |
| AAGGCCGCTGATACCGATAGTTGCGACGTTAG |
| CCCAGCAGGCGAAAAATCCCTTATAAATCAAGCCGGCG |
| TAAATCAAAATAATTCGCGTCTCGGAAACCAGGCAAAGGGAAGG |
| GAGACAGCTAGCTGATAAATTAATTTTTGT |
| TTTGGGGATAGTAGTAGCATTAAAAGGCCG |
| GCTTCAATCAGGATTAGAGAGTTATTTTCA |
| CGTTTACCAGACGACAAAGAAGTTTTGCCATAATTCGA |
| TGACAACTCGCTGAGGCTTGCATTATACCAAGCGCGATGATAAA |
| TCTAAAGTTTTGTCGTCTTTCCAGCCGACAA |
| TCAATATCGAACCTCAAATATCAATTCCGAAA |
| GCAATTCACATATTCCTGATTATCAAAGTGTA |
| AGAAAACAAAGAAGATGATGAAACAGGCTGCG |
| ATCGCAAGTATGTAAATGCTGATGATAGGAAC |
| GTAATAAGTTAGGCAGAGGCATTTATGATATT |
| CCAATAGCTCATCGTAGGAATCATGGCATCAA |
| AGAGAGAAAAAAATGAAAATAGCAAGCAAACT |
| TTATTACGAAGAACTGGCATGATTGCGAGAGG |

| |
|---|
| GCAAGGCCTCACCAGTAGCACCATGGGCTTGA |
| TTGACAGGCCACCACCAGAGCCGCGATTTGTA |
| TTAGGATTGGCTGAGACTCCTCAATAACCGAT |
| TCCACAGACAGCCCTCATAGTTAGCGTAACGA |
| AACGTGGCGAGAAAGGAAGGGAAACCAGTAA **TTA TAC ATC TAT TTC TTC ATT ATT CAC TTA CTA** |
| TCGGCAAATCCTGTTTGATGGTGGACCCTCAA |
| AAGCCTGGTACGAGCCGGAAGCATAGATGATG |
| CAACTGTTGCGCCATTCGCCATTCAAACATCA **TTT TAG GTA AAT T TTG ATT GTG AGG AAG** |
| GCCATCAAGCTCATTTTTTAACCACAAATCCA |
| CAACCGTTTCAAATCACCATCAATTCGAGCCA |
| TTCTACTACGCGAGCTGAAAAGGTTACCGCGC |
| CCAACAGGAGCGAACCAGACCGGAGCCTTTAC |
| CTTTTGCAGATAAAAACCAAAATAAAGACTCC **TTT TAG GTA AAT T TTG ATT GTG AGG AAG** |
| GATGGTTTGAACGAGTAGTAAATTTACCATTA |
| TCATCGCCAACAAAGTACAACGGACGCCAGCA |
| ATATTCGGAACCATCGCCCACGCAGAGAAGGA **TTA TAC ATC TAT TTC TTC ATT ATT CAC TTA CTA** |
| TAAAAGGGACATTCTGGCCAACAAAGCATC |
| ACCTTGCTTGGTCAGTTGGCAAAGAGCGGA |
| ATTATCATTCAATATAATCCTGACAATTAC |
| CTGAGCAAAAATTAATTACATTTTGGGTTA |
| TATAACTAACAAAGAACGCGAGAACGCCAA |
| CATGTAATAGAATATAAAGTACCAAGCCGT |
| TTTTATTTAAGCAAATCAGATATTTTTTGT |
| TTAACGTCTAACATAAAAACAGGTAACGGA |
| ATACCCAACAGTATGTTAGCAAATTAGAGC |
| CAGCAAAAGGAAACGTCACCAATGAGCCGC |
| CACCAGAAAGGTTGAGGCAGGTCATGAAAG |
| TATTAAGAAGCGGGGTTTTGCTCGTAGCAT |
| TCAACAGTTGAAAGGAGCAAATGAAAAATCTAGAGATAGA |
| TCAAATATAACCTCCGGCTTAGGTAACAATTTCATTTGAAGGCGAATT |
| GTAAAGTAATCGCCATATTTAACAAAACTTTT |
| TATCCGGTCTCATCGAGAACAAGCGACAAAAG |
| TTAGACGGCCAAATAAGAAACGATAGAAGGCT |
| CGTAGAAAATACATACCGAGGAAACGCAATAAGAAGCGCA |
| GCGGATAACCTATTATTCTGAAACAGACGATTGGCCTTGAAGAGCCAC |
| TCACCAGTACAAACTACAACGCCTAGTACCAG |
| ACCCTTCTGACCTGAAAGCGTAAGACGCTGAG |
| AGCCAGCAATTGAGGAAGGTTATCATCATTTT |
| GCGGAACATCTGAATAATGGAAGGTACAAAAT |
| CGCGCAGATTACCTTTTTTAATGGGAGAGACT |
| ACCTTTTTATTTTAGTTAATTTCATAGGGCTT |
| AATTGAGAATTCTGTCCAGACGACTAAACCAA |

| |
|---|
| GTACCGCAATTCTAAGAACGCGAGTATTATTT |
| ATCCCAATGAGAATTAACTGAACAGTTACCAG |
| AAGGAAACATAAAGGTGGCAACATTATCACCG |
| TCACCGACGCACCGTAATCAGTAGCAGAACCG |
| CCACCCTCTATTCACAAACAAATACCTGCCTA |
| TTTCGGAAGTGCCGTCGAGAGGGTGAGTTTCG |
| CTTTAGGGCCTGCAACAGTGCCAATACGTG |
| CTACCATAGTTTGAGTAACATTTAAAATAT |
| CATAAATCTTTGAATACCAAGTGTTAGAAC |
| CCTAAATCAAAATCATAGGTCTAAACAGTA |
| ACAACATGCCAACGCTCAACAGTCTTCTGA |
| GCGAACCTCCAAGAACGGGTATGACAATAA |
| AAAGTCACAAAATAAACAGCCAGCGTTTTA |
| AACGCAAAGATAGCCGAACAAACCCTGAAC |
| TCAAGTTTCATTAAAGGTGAATATAAAAGA |
| TTAAAGCCAGAGCCGCCACCCTCGACAGAA |
| GTATAGCAAACAGTTAATGCCCAATCCTCA |
| AGGAACCCATGTACCGTAACACTTGATATAA |
| GCACAGACAATATTTTTGAATGGGGTCAGTA |
| TTAACACCAGCACTAACAACTAATCGTTATTA |
| ATTTTAAAATCAAAATTATTTGCACGGATTCG |
| CCTGATTGCAATATATGTGAGTGATCAATAGT |
| GAATTTATTTAATGGTTTGAAATATTCTTACC |
| AGTATAAAGTTCAGCTAATGCAGATGTCTTTC |
| CTTATCATTCCCGACTTGCGGGAGCCTAATTT |
| GCCAGTTAGAGGGTAATTGAGCGCTTTAAGAA |
| AAGTAAGCAGACACCACGGAATAATATTGACG |
| GAAATTATTGCCTTTAGCGTCAGACCGGAACC |
| GCCTCCCTCAGAATGGAAAGCGCAGTAACAGT |
| GCCCGTATCCGGAATAGGTGTATCAGCCCAAT |
| AGATTAGAGCCGTCAAAAAACAGAGGTGAGGCCTATTAGT |
| GTGATAAAAAGACGCTGAGAAGAGATAACCTTGCTTCTGTTCGGGAGA |
| GTTTATCAATATGCGTTATACAAACCGACCGT |
| GCCTTAAACCAATCAATAATCGGCACGCGCCT |
| GAGAGATAGAGCGTCTTTCCAGAGGTTTTGAA |
| GTTTATTTTGTCACAATCTTACCGAAGCCCTTTAATATCA |
| CAGGAGGTGGGGTCAGTGCCTTGAGTCTCTGAATTTACCGGGAACCAG |
| CCACCCTCATTTTCAGGGATAGCAACCGTACT |
| CTTTAATGCGCGAACTGATAGCCCCACCAG **TTA TAC ATC TAT TTC TTC ATT ATT CAC TTA CTA** |
| CAGAAGATTAGATAATACATTTGTCGACAA |
| CTCGTATTAGAAATTGCGTAGATACAGTAC |
| CTTTTACAAAATCGTCGCTATTAGCGATAG **TTT TAG GTA AAT T TTG ATT GTG AGG AAG** |
| CTTAGATTTAAGGCGTTAAATAAAGCCTGT |
| TTAGTATCACAATAGATAAGTCCACGAGCA |

| |
|---|
| TGTAGAAATCAAGATTAGTTGCTCTTACCA |
| ACGCTAACACCCACAAGAATTGAAAATAGC |
| AATAGCTATCAATAGAAAATTCAACATTCA **TTT TAG GTA AAT T TTG ATT GTG AGG AAG** |
| ACCGATTGTCGGCATTTTCGGTCATAATCA |
| AAATCACCTTCCAGTAAGCGTCAGTAATAA |
| GTTTTAACTTAGTACCGCCACCCAGAGCCA **TTA TAC ATC TAT TTC TTC ATT ATT CAC TTA CTA** |

**Supplementary Note 1: Quantitative principles of Repeat DNA-PAINT**

**Setting imager concentration in DNA-PAINT**

In a DNA-PAINT experiment one typically has a given density $\rho_{DS}$ of docking strands per unit area in the sample, proportional to the density of target epitopes. For super-resolution imaging, only a small fraction $f_{DS}$ of these strands should be occupied by imagers at any time, typically $< 5\%$, so that the event density per unit area $E = \rho_{DS} f_{DS}$ is below a maximal value $E_{max}$, defined by the requirement that the probability of two binding events occurring simultaneously in a diffraction-limited volume remains small, *i.e.*

$$E_{max} = \rho_{DS} f_{DS,max} \,. \qquad \text{(Eq S1)}$$

In experiments, one often quantifies event rates, i.e. the number of events in a certain image region that occur per frame. The event density $E$ described here and the experimentally measured event rate are proportional (by the factor of the area of the image region) and therefore event rates are often used synonymously for the event densities discussed here, as in the main text of this work.

To minimize the total image-acquisition time, the fraction $f_{DS}$ should be adjusted so that the event density is as close as possible to the maximal value: $E \approx E_{max}$. Since $f_{DS} \ll 1$, the fraction of occupied docking sites scales linearly with imager concentration $[I]$

$$f_{DS} = \frac{[I]}{K_d}, \qquad \text{(Eq S2)}$$

where the dissociation constant $K_d$ depends on the docking domain/imager strand design.

One should therefore seek to adjust imager concentration $[I]$ to achieve

$$E_{max} \approx \rho_{DS} \cdot \frac{[I]}{K_d} \,. \qquad \text{(Eq S3)}$$

We can thus define the optimal imager concentration $[I]_0 = E_{max} K_d / \rho_{DS}$.

Equations (S1-S3) also show that should one chose to reduce $[I]$ to suppress free-imager background (we show the effect of $[I]$ on background in **Fig. 2b**) and non-specific events (we quantify the effect of $[I]$ on non-specific imager binding in **Fig. 3a**) then the event density would also be reduced to $E \ll E_{max}$. Such a reduction to non-optimal event densities is problematic as it proportionally increases image-acquisition timescales, which is often incompatible with the limitations imposed by sample degradation and mechanical stability of the imaging setups.

**Domain repeats: Reducing imager concentration**

With Repeat DNA-PAINT, by introducing multiple imager-binding domains, we increase the effective density of docking sites in direct proportion to the domain repeat number $N$ (as we show in **Fig. 1**). In other words, with domain repeats the effective docking site density is now

$$\rho_{DS,eff} = \rho_{DS} \cdot N \,. \qquad \text{(Eq S4)}$$

Thus, to get the same $E_{max}$ we can reduce the optimal imager concentration by the factor $N$,

$$E_{max} = \rho_{DS,eff} \cdot \frac{[I]_R}{K_d} = \rho_{DS} \cdot N \cdot \frac{[I]_0}{N} \cdot \frac{1}{K_d} = \rho_{DS} \cdot \frac{[I]_0}{K_d}, \qquad \text{(Eq S5)}$$

where $[I]_R = [I]_0 / N$ is the Repeat DNA-PAINT optimal imager concentration. In the main text (**Figs. 2 and 3**) we demonstrate the benefits of a lower imaging concentration on free-imager backgrounds and non-specific imager binding.

**Domain repeats: Accelerating DNA-PAINT data acquisition**

In a DNA-PAINT experiment the single-frame integration time should be of the same order of magnitude as the duration of an individual docking-imager binding event. Therefore, an acceleration in data acquisition, which requires an increase in frame-rate, must be accompanied by a proportional shortening of binding events. The latter can be achieved by using imagers which display lower affinity for the docking domains, corresponding to higher off-rates and dissociation constants $K_d$. Thus, from Eq. (S3), to achieve a similar $E_{max}$, the imager concentration needs to increase in direct proportion to the acceleration required. This rapidly becomes prohibitive in terms of background photon levels and non-specific imager binding, both of which increase with $[I]$.

For example, to accelerate acquisition by a factor $N$, the dissociation constant should be increased to $K_{d,acc} = K_d \cdot N$, which in conventional DNA-PAINT would require a much higher imager concentration $[I]_{acc} = N[I]_0$ to maintain event densities at the maximal spatial density $E_{max}$ . This would in turn greatly increase background fluorescence and non-specific binding. Instead, with Repeat DNA-PAINT we achieve $E_{max}$ with the same imager concentration $[I]_0$ as un-accelerated conventional DNA-PAINT (that uses imagers with the larger dissociation constant $K_d$):

$$E_{max} = \rho_{DS,eff} \cdot \frac{[I]_0}{K_{d,acc}} = \rho_{DS} \cdot N \cdot [I]_0 \frac{1}{K_d/N} = \rho_{DS} \cdot \frac{[I]_0}{K_d} \ . \qquad \text{(Eq S6)}$$

Note also that Eqs (S1-S6) provide a framework to design strategies which compromise between the key benefits afforded by Repeat DNA-PAINT: accelerating acquisition as well as reducing backgrounds and non-specific events.

**Supplementary Note 2. Estimation of the thermal relaxation timescale of docking motifs.**

Here we show that the timescales of thermal fluctuations of the docking motifs are very fast compared to the rates of photon emission during a binding event. Hence, the physical locations from which photons are emitted are uncorrelated and drawn from the equilibrium distribution of fluorophore positions we quantify numerically in **Fig. 5a**. Since the latter are symmetric around the anchoring location of the docking motif, Repeat DNA-PAINT does not introduce any random bias in localization, and the net effect of physically extending the docking motifs is simply that of negligibly widening the camera image of a blink (**Fig. 5b**).

To estimate the timescales of thermal relaxation of the docking-imager complexes we chose to neglect the presence of the imager (which we justify below) and regard the docking strand as a flexible bead chain, whose dynamics can be described through the Rouse or Zimm models.[1]

In the Rouse model, hydrodynamic interactions are ignored, and the thermal decorrelation timescale of each normal (Fourier) mode is given by

$$\tau_{pR} = \frac{2N^2a^2b\eta}{(\pi k_B T p^2)}. \qquad \text{(Eq S7)}$$

In the Zimm model, hydrodynamic interactions are considered, and decorrelation timescales are given by

$$\tau_{pZ} = \frac{(\sqrt{N}a)^3}{\sqrt{(3\pi p^3)}}\frac{\eta}{k_B T}. \qquad \text{(Eq S8)}$$

In Eqs (S7) and (S8), $N$ is the number of beads in the polymer, $b$ the bead radius, $a$ is the bead-to-bead distance, $\eta$ the dynamic viscosity of the fluid (water), and $p$ is the Fourier mode number, with $p = 1$ being the slowest decaying mode. We chose $b = 4$ nm and $b = 9$, as parametrized from Fluorescence Correlation spectroscopy data recorded for ssDNA.[1]

The docking motif that stretches the furthest from the anchoring point, and thus the one with the slowest fluctuation modes, is 6xRD (**Fig. 1a**). Note that although 10xRD is overall longer, it is tethered through the middle and the two dangling segments are individually shorter than 6xRD. The contour length of 6xRD is 56 nt, corresponding to[2] 38 nm and $N = 4$, using the empirically derived values of $a$ and $b$. At room temperature (22 °C) and using $\eta = 1$ mPa s, the Rouse model (Eq. S7) estimates a maximum decay time ($p = 1$) $\tau_{1R} \approx 500$ ns, while the Zimm model (Eq. S8) produces $\tau_{1Z} \approx 800$ ns.

The mean interval between subsequent photon emissions can be estimated by the ratio of the binding time (~500 ms) to the dye photon budget (<10,000), rendering ~50 µs. This value is at least two orders of magnitude greater than $\tau_{1R}$ and $\tau_{1Z}$, ensuring that the docking motif can fully sample its configurational space between subsequent photon emissions. The single-frame integration time (>10 ms) is even more comfortably in excess of the docking-motif decorrelation timescales. Note that in this calculation we neglect the presence of the imager hybridized to docking motif, which results in a stiffer dsDNA segment that could influence the polymer properties of the docking strand. Given the vast difference between the estimated equilibration and photon-emission timescales, however, we argue that this approximation has a negligible effect with respect to our conclusion.
